# Spatial structures of fungal DNA assemblages revealed with eDNA metabarcoding in a forest river network in western Japan

**DOI:** 10.1101/637686

**Authors:** Shunsuke Matsuoka, Yoriko Sugiyama, Hirotoshi Sato, Izumi Katano, Ken Harada, Hideyuki Doi

## Abstract

Growing evidence has revealed high diversity and spatial heterogeneity of fungal communities including in local habitats in terrestrial ecosystems. These findings highlight the considerable sampling effort, analysis time, and costs required for the investigation of fungal diversity over large spatial scales. Recently, the analysis of environmental DNA in river water has been undertaken to study the biodiversity of organisms, such as animals and plants, in both aquatic and terrestrial habitats. However, previous studies have not investigated the spatial structure of fungal DNA assemblages in river water. In the present study, we investigate fungal DNA assemblages and their spatial structure using environmental DNA metabarcoding in water across different branches of river over forest landscapes. The river water was found to contain both phylogenetically and functionally diverse fungal DNA, including aquatic and terrestrial fungi, such as plant decomposers and mycorrhizal fungi. These fungal DNA assemblages were more similar within, rather than between, branches. In addition, the assemblages were more similar between spatially closer branches. These results imply that information on the terrestrial and aquatic fungal compositions of watersheds, and therefore their spatial structure can be obtained by investigating the fungal DNA assemblages in river water.

## Introduction

Information on biodiversity composition is essential to understand the ecological processes and functions of ecosystems. Understanding patterns of biodiversity is fundamental for the maintenance of ecological processes during this era of environmental change (Margules and Pressey, 2000; Pecl et al., 2017). The Kingdom Fungi boasts high species diversity and undertakes unique functions in both terrestrial and aquatic ecosystems via the decomposition of organic substrates and symbiotic or parasitic interactions with other organisms, such as animals and plants (Peay et al., 2016; Hawksworth and Lücking, 2017; Grossart et al., 2019). Recently, with the growing use of DNA metabarcoding techniques, an increasing number of studies are investigating fungal diversity on a variety of substrates in terrestrial and aquatic habitats (Duarte et al., 2015; Peay et al., 2016; Nilsson et al., 2019). However, as fungi show high local diversity and spatial heterogeneity (Bahram et al., 2015, 2016), the investigation of fungal diversity over large spatial scales, such as landscape (tens of km) and regional scales (hundreds to thousands of km) requires a significant amount of sampling effort, time for analysis, and costs.

Recently, the availability of environmental DNA in river water for biodiversity exploration has been discussed in several studies (Deiner et al., 2016; Nakagawa et al. 2018). Deiner et al. (2016) suggest that river water can integrate DNA from the catchment area (~tens of km) and work as conveyer belts of diversity information for various eukaryotes in both aquatic and terrestrial habitats. It is expected that the spores and mycelia of terrestrial fungi enter the rivers allowing for the detection of the terrestrial as well as the aquatic fungi (Voronin, 2014). Indeed, recent studies have detected the DNA of supposedly terrestrial fungi from river water (Deiner et al., 2016; LeBrun et al., 2018), indicating that the diversity information of fungi in a catchment area can be accessed by examining fungal DNA assemblages in river water. In this scenario, even those not familiar with the identification of fungi could infer the fungal diversity of a catchment area from river water samples. By using this method, the fungal diversity information of any habitat type can be obtained in less time and at a lower cost than through traditional methods.

Rivers typically consist of a network of multiple branches that originate from the main stream. Terrestrial fungal communities have spatial structures wherein geographically closer sites share similar fungal communities which reflects the surrounding environment and dispersal ability of each species (Talbot et al., 2014; Matsuoka et al., 2016b; Peay et al., 2016). Assuming that fungal DNA assemblages in river water reflect the fungal community compositions of surrounding land, the assemblages also indicate the spatial structure. Rivers have a directional water flow, from upstream to downstream. Therefore, the DNA assemblages of river water could also show river-specific structures. Samples from the same branch may be more similar to one another than those from different branches. Information regarding fungal DNA assemblages in river water and their spatial structure is essential for investigations into the broad spatial diversity of fungi. However, no previous studies have investigated fungal DNA assemblages in river water focusing on the spatial structure.

In the present study, we aim to investigate fungal DNA assemblages in river water across branches over forest landscapes and clarify its spatial structure. In particular, we investigate (1) whether fungal DNA assemblages are more similar within, rather than between, river branches, and (2) whether fungal DNA assemblages show a spatial structure that reflects geographical distance or environmental factors.

## Methods

### Study sites and sampling

In the present study, the same water samples as Katano et al. (2017) were used. Study sites are located in the headwater tributaries of the Ibo River in Hyogo, Japan (Fig. 1). Seven sites are located in the Akazai River (A1-A7) and twelve sites in the tributaries of the Ibo River (I1-I12). The orders of the streams and width of the tributaries are 1–2 and 3–5 m, respectively. A1-A7, I7 and I9, and I8 and I9 are located on the same tributary. At each sampling site, longitude, latitude, and altitude were recorded using a GPS device (Germin etrex 30x).

**Fig. 1.**
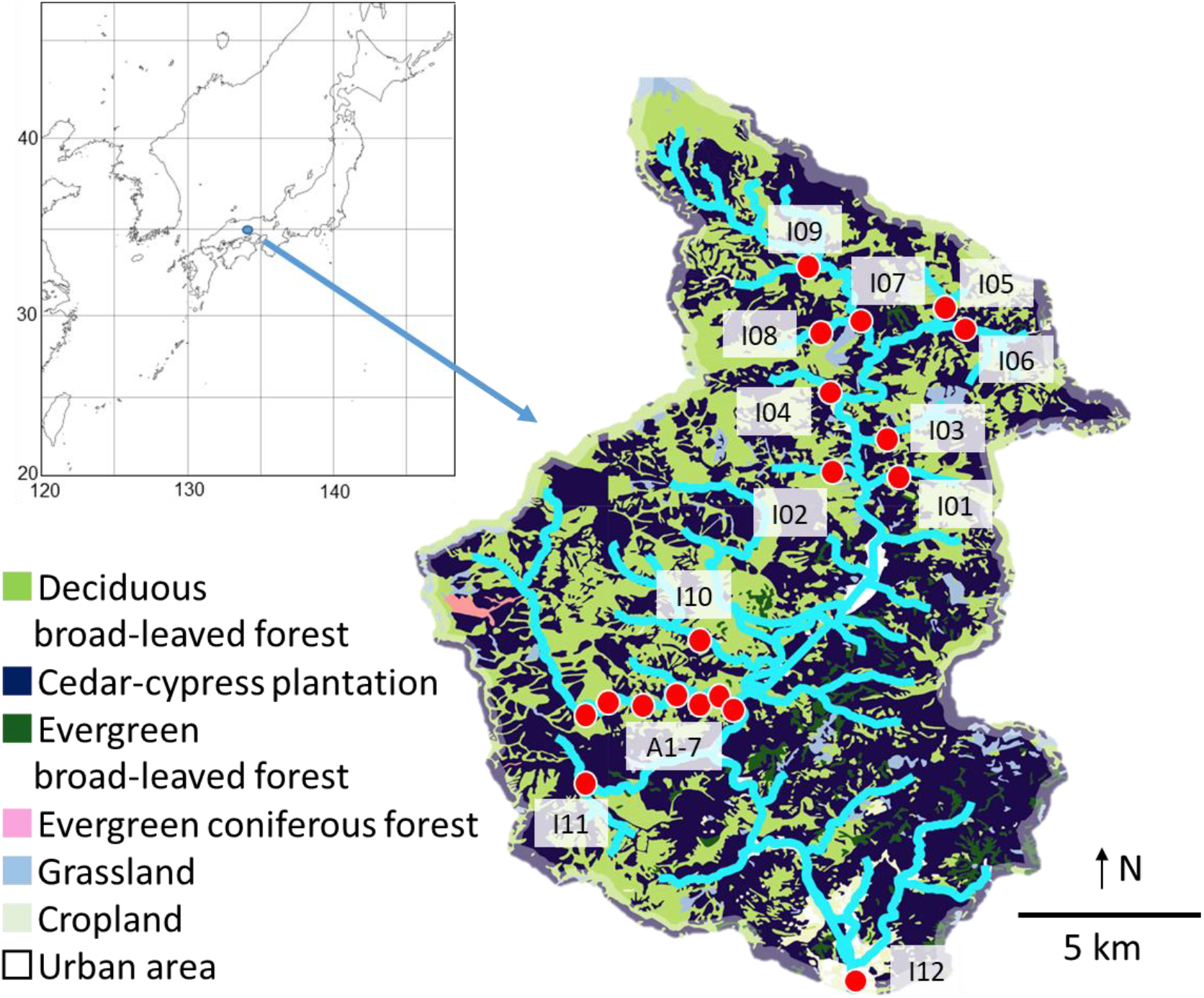
Sampling sites and land use of the catchment area.

A 1 L sample was collected from the surface of the water at the center of each stream in the study sites using a DNA-free polypropylene bottle. Samples were stored in a cooler box with a blank. The blank contained 1 L distilled water which was brought to the field and treated identically to the other water sampling bottles, except that it was not opened at the field sites. The sampling was conducted in late September, 2015. The collected water samples and blank were vacuum-filtered through 47 mm GF/F glass filters (pore size 0.7 μm, GE Healthcare, Little Chalfont, UK). The filter was then wrapped in commercial aluminum foil and stored at −20°C before DNA extraction.

### Molecular Experiments

The methods described by Uchii et al. (2016) were followed to extract DNA from the filters. Each filter was incubated in 400 μl of Buffer AL (Qiagen, Hilden, Germany) and 40 μl of Proteinase K (Qiagen, Hilden, Germany), using a Salivette® tube (Sarstedt, Nümbrecht, Germany) at 56°C for 30 min. The Salivette® tube with the filter was centrifuged at 3,500 × g for 5 min. Next, 220 μl of TE buffer (10 mM Tris-HCl and 1 mM EDTA, pH: 8.0) was added to the filter and centrifuged at 5,000 × g for 5 min. The dissolved DNA in the eluted solution was purified using a DNeasy Blood & Tissue Kit (Qiagen, Hilden, Germany) according to the manufacturer’s protocol. The final volume of the extracted sample was eluted in 100 μl of Buffer AE of the DNeasy Blood & Tissue Kit.

For MiSeq sequencing, the fungal internal transcribed spacer 1 (ITS 1) region of rDNA was amplified. The ITS region has been proposed as the formal fungal barcode (Schoch et al., 2012). The methods of DNA analysis followed those described by Ushio et al. (2017) with some modifications, such as number of PCR cycles. The first-round PCR (first PCR) amplified the ITS1 region using the ITS1-F-KYO2 and ITS2-KYO2 primer set, which is capable of amplifying the ITS1 region of most fungal groups (Toju et al., 2012). An Illumina sequencing primer and six random bases (N) were combined to produce each primer. Thus, the forward primer sequence was: 5′-. *ACA CTC TTT CCC TAC ACG ACG CTC TTC CGA TCT* NNNNNN TAG AGG AAG TAA AAG TCG TAA −3′ and the reverse primer sequence was: 5′-*GTG ACT GGA GTT CAG ACG TGT GCT CTT CCG ATC T* NNNNNN TTY RCT RCG TTC TTC ATC-3′. The italic and normal letters represent MiSeq sequencing primers and fungi-specific primers, respectively. The six random bases (N) were used to enhance cluster separation on the flowcells during initial base call calibrations on MiSeq.

The 1^st^ PCR was performed in a 12 μl volume with the buffer system of KODFX NEO (TOYOBO, Osaka, Japan), which contained 2.0 μl of template DNA, 0.2 μl of KOD FX NEO, 6.0 μl of 2× buffer, 2.4 μl of dNTP, and 0.7 μl each of the two primers (5 μM). The PCR conditions were as follows; an initial incubation for 2 min at 94°C followed by 5 cycles of 10 s at 98°C, 30 s at 68°C for annealing and 30 s at 68°C, 5 cycles of 10 s at 98°C, 30 s at 65°C and 30 s at 68°C; 5 cycles of 10 s at 98°C, 30 s at 62°C and 30 s at 68°C; 25 cycles of 10 s at 98°C, 30 s at 59°C and 30 s at 68°C, and a final extension of 5 min at 68°C. Eight replicate first-PCRs (per sample) were performed to mitigate the reaction-level PCR bias. Then, the duplicated first PCR amplicons (per sample) were combined, resulting in a template per sample for the second PCR. The PCR templates were purified using Agencourt AMPure XP (PCR product: AMPure XP beads = 1:0.8; Beckman Coulter, Brea, California, USA) before the second PCR.

The second PCR amplified the first PCR amplicons using the primers (forward) 5′-*AAT GAT ACG GCG ACC ACC GAG ATC TAC AC* XXXXXXXX TCG TCG GCA GCG TCA GAT GTG TAT AAG AGA CAG-3′ and (reverse) 5′-*CAA GCA GAA GAC GGC ATA CGA GAT* XXXXXXXX GTC TCG TGG GCT CGG AGA TGT GTA TAA GAG ACA G-3′. The italic and normal letters represent the MiSeqP5/P7 adapter and sequencing primers, respectively. The 8X bases represent dual-index sequences inserted to identify different samples (Hamady et al., 2008). The second PCR was carried out with 12 μl reaction volume containing 1.0 μl of template, 6 μl of 2× KAPA HiFi HotStart ReadyMix (KAPA Biosystems, Wilmington, Washington, USA), 1.4 μl of each primer (2.5 μM), and 2.2 μl of sterilized distilled water. The PCR conditions were as follows; an initial incubation for 3 min at 95°C followed by 12 cycles of 20 s at 98°C, 15 s at 72°C for annealing and extension, and a final extension of 5 min at 72°C.

The indexed second PCR amplicons were pooled to make a library to be sequenced on MiSeq. The volume of each sample added to the library was adjusted to normalize the concentrations of each second PCR product. The pooled library was purified using Agencourt AMPure XP. A target-sized DNA of the purified library (approximately 380–510 base pairs [bp]) was then excised using E-Gel SizeSelect (ThermoFisher Scientific, Waltham, MA, USA). The double-stranded DNA concentration of the library was then adjusted to 4 nmol/L using Milli-Q water, and the DNA sample was applied to the Illumina MiSeq platform at Ryukoku University, Japan. The sequence data were deposited in the Sequence Read Archive of the DNA Data Bank of Japan (accession number: DRA007786).

### Bioinformatics

The procedures used for bioinformatics and data analyses followed those described previously (Matsuoka et al., 2016a; Ushio et al., 2017). The raw MiSeq data were converted into FASTQ files using the bcl2fastq program provided by Illumina. The FASTQ files were then demultiplexed using the commands implemented in Claident pipeline (Tanabe and Toju, 2013; software available online: https://www.claident.org/). This process was adopted rather than using FASTQ files demultiplexed by the Illumina MiSeq default program to remove sequences whose 8-mer index positions included nucleotides with low quality scores (i.e., Phred score < 30). The forward and reverse sequences were then merged with each other. The total 611,092 reads (32162 ± 10784 reads per sample, mean ± SD, n = 19) were assembled using Claident v0.2.2016.07.05. First, short (<150 bp) reads were removed, then potentially chimeric sequences and pyrosequencing errors were removed using UCHIME v4.2.40 (Edger et al., 2011) and algorithms in CD-HIT-OTU (Li et al., 2012), respectively. The remaining sequences were assembled at a threshold similarity of 97% (Osono, 2014), and the resulting consensus sequences represented molecular operational taxonomic units (OTUs). Then, singletons were removed. Through these procedures, 2,549 OTUs (602,304 reads) in total were obtained. An OTU table (i.e., matrix of OTUs and samples with sequence reads in each cell entry) was generated after this process, and subsequent statistical analysis was performed as described below in the statistics section. For each of the obtained OTUs, taxonomic identification was conducted based on the query-centric auto-k-nearest-neighbor (QCauto) method (Tanabe and Toju, 2013) and subsequent taxonomic assignment with the lowest common ancestor algorithm (Huson et al., 2007) using Claident. The functional guild of each fungal OTU was estimated based on the FUNGuild database (Nguyen et al., 2016).

### Vegetation information of catchment area

By using QGIS ver.2.14.8, information on the gross area and vegetation of the catchment area of each sampling site was calculated. First, the catchment area of each site was calculated based on the GPS coordinates of the site. Then, by consulting the vegetation map made by the Ministry of the Environment of Japan, vegetation of the catchment area was divided into eight categories (deciduous broad-leaved forest, cedar-cypress plantation, ever-green broad-leaved forest, ever-green coniferous forest, grassland, cropland, and urban area) and the area of each category was calculated (Fig. 1).

### Statistical analyses

The OTU table (i.e., fungal OTUs × samples matrix, in which a cell entry indicates the number of sequence reads of each OTU in each sample) was used for the following analysis. First, coverage-based rarefaction was conducted to standardize the effect of sequencing depth (Chao and Jost, 2012, rarefaction curves for each sample are shown in Fig. S1). Then, 39 OTUs (718 reads) that were identified as non-fungal organisms were discarded and the remaining 1,956 OTUs (175,101 reads) were used in further analysis (Table S1). The presence or absence of the OTUs was recorded for each sample regardless of the number of reads and used for all statistical analyses as binary data. All analyses were performed using R v.3.4.3 (R Core Team 2017). Note that in the present study, analyses described below were conducted on the datasets with unidentified OTUs. However, to investigate the effect of inclusion of unidentified OTUs, and the use of presence/absence data instead of read number, additional analyses were conducted by using datasets for only fungal OTUs (1,031 OTUs, see Table S1), and by considering read numbers of each OTU. The results for these additional analyses are provided in the Appendix, but the results did not differ among these analyses.

The variation of community composition of OTUs among sampling sites and the community dissimilarity of the OTUs among plots were visualized using non-metric multidimensional scaling (NMDS). Presence/absence data of OTUs for each site were converted into a dissimilarity matrix using the standardized effect size (SES) of the Jaccard dissimilarity index to show the community dissimilarity without dependence on richness gradient among sites, which is an issue of long-standing importance (Chase and Myers, 2011). The SES was defined as: (β_obs_ -β_null_) / β_sd_, where β_obs_ is the observed β diversity, β_null_ is the mean of the null distribution of β diversity, and β_sd_ is the standard deviation of the null distribution. The null distribution was calculated based on 999 randomizations preserving both the site occurrence and the OTU richness. The SES value indicates the magnitude of the deviation from the expectation of a random assembly process, and positive and negative values indicate more and less β-diversity, respectively, than expected by chance. I12 is the only site located in an urban area, and the NMDS ordination indicates that the community in I12 is largely different from the others (Fig. S2). Therefore, we excluded this site from further analyses. The correlation of NMDS structure with geographic coordinates (latitude and longitude) was tested through permutation tests (‘envfit’ command in the vegan package, 9999 permutations) to show the spatial structure of OTU composition among sites. Further, only the results for presence/absence data are shown. Then, to test whether the OTU compositions are more similar within a tributary than between tributaries, SES values were compared calculating 99% confidence interval (CI).

To test the effects of environmental variables (i.e., elevation, catchment area, and vegetation) and spatial distance on the OTU composition, a multivariate permutational analysis of variance (PERMANOVA) was implemented (‘adonis’ command in the vegan package, 9999 permutations). In these analyses, only one sample was used for each branch as samples from the same branch (A1-7 and I7-8) share most of their catchment. Therefore, eleven samples (A4, I1-6, and I8-11) were used for the following analyses and others were excluded. Preliminary analyses confirmed that the choice of sample inside the branch does not affect the results (data were used in Table S1). Presence/absence data of ECM fungal OTUs for each plot were converted into a dissimilarity matrix using the modified Raup–Crick dissimilarity index (Chase et al., 2011). The Raup–Crick dissimilarity index is calculated based on the null model approach, which is akin to the SES value described above. Then, vegetation and spatial variables were calculated. Vegetation variables were extracted based on principal coordinate analysis (PCA). First, the vegetation data of the catchment area of each sampling site were transferred into the Euclidian distance matrix. Then, PCA vectors were extracted. The first two PCA axes explained the variations in the vegetation among sites 86.9% and 11.2%, respectively, (Fig. S3) and were used for the analysis. Spatial variables were extracted based on principal components of neighbor matrices (PCNM, Borcard et al., 2004). The PCNM analysis produced a set of orthogonal variables derived from the geographical coordinates of the locations of sampling sites. Two PCNM variables that best accounted for autocorrelation were used for the analysis.

## Results

### Taxonomic assignment

In total, the filtered 175,101 reads from 19 samples were grouped into 1,956 OTUs with 97% sequence similarity (Table S1). In total, 770 OTUs were assigned as Ascomycota (38.4 % of the total number of fungal OTUs), 177 OTUs as Basidiomycota (8.8 %), and 38 OTUs as Chytridiomycota (1.9 %). The remaining 941 OTUs were not assigned to any phylum. The proportions of OTU numbers of each phylum were similar between the sampling sites except for I12 (Fig. S4). FUNGuild assigned 539 OTUs to functional guilds, 147 OTUs of which were saprotrophs and the others included pathogens (117 OTUs) and symbionts (19 OTUs) (Table S.1). The detected OTU included aquatic hyphomycetes, e.g., *Flagellospora* sp. (OTU_0413) and *Tetracladium marchalianum* (OTU_1491), were dominantly detected from 19 and 16 of 19 samples, respectively. On the other hand, OTUs belonging to terrestrial taxonomic or functional groups were also detected; plant decomposer (e.g., *Mycena* sp. (OTU_0020)); plant pathogen (e.g., *Ciboria shiraiana* (OTU_1523)); and mycorrhizal fungi (e.g., Glomeromycetes sp. (OTU_1449) and *Tuber* sp. (OTU_0010)).

### Spatial structures of fungal OTU composition

The Mantel test and NMDS ordination revealed the biogeographic changes in the fungal OTU compositions. The dissimilarity of OTU composition among samples was significantly correlated with distances of latitude and longitude (Mantel test, latitude, *r* = 0.585, P < 0.001; longitude, *r* = 0.510, P < 0.001). The NMDS ordination showed the geographical change of OTU composition among sites (Fig 2, stress value = 0.154). The ordination was significantly correlated with the latitude and longitude (‘envfit’ function; latitude, r^2^ = 0.737, P < 0.001; longitude, r^2^ = 0.781, P < 0.001). The dissimilarity indices of OTU composition (SES value of Jaccard index) were significantly lower than 0 when compared between the sites in the same branch, higher when compared between distant branches, and intermediate when compared between adjacent branches (Fig. 3, based on the 99% confidence interval). This indicates that OTU compositions are more significantly similar inside the branch than expected by chance and more different between spatially distinct different branches. PERMANOVA showed that the differences in OTU composition among the branches were only related to the spatial variable (PCNM1) (F-model = 6.938, R^2^ = 0.432, P = 0.009) and did not significantly relate with the other environmental variables including elevation and vegetation PCA axis (Table 1).

**Table 1.** PERMANOVA results for the composition of DNA assemblages

|  | Degree of<br>freedom | Sums of<br>Squares | F Model | R<sup>2</sup> | P value |
| --- | --- | --- | --- | --- | --- |
| null | 10 | 1.801 |  | 1 |  |
| Elevation | 1 | 0.392 | 2.502 | 0.218 | 0.169 |
| Catchment area | 1 | 0.248 | 1.434 | 0.137 | 0.356 |
| Vegetation PC1 | 1 | 0.150 | 0.819 | 0.083 | 0.509 |
| Vegetation PC2 | 1 | 0.001 | 0.003 | 0.001 | 0.838 |
| PCNM1 | 1 | 0.778 | 6.838 | 0.432 | 0.009 |
| PCNM2 | 1 | -0.125 | -0.584 | -0.069 | 0.942 |

**Fig. 2.**
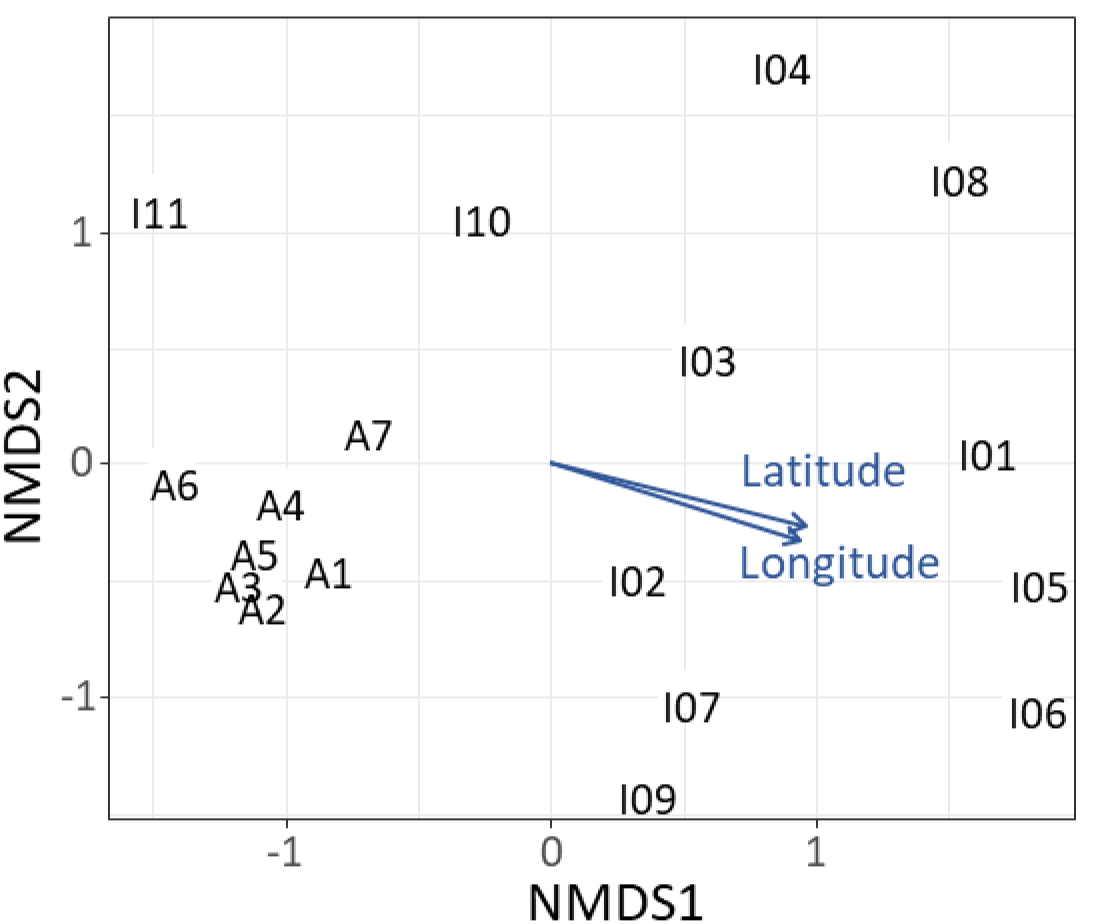
Dissimilarity of the DNA assemblages among sites as revealed by nonmetric multidimensional scaling (NMDS) ordination (stress value = 0.154). Numbers are consistent with site numbers in Fig. 1.

**Fig. 3.**
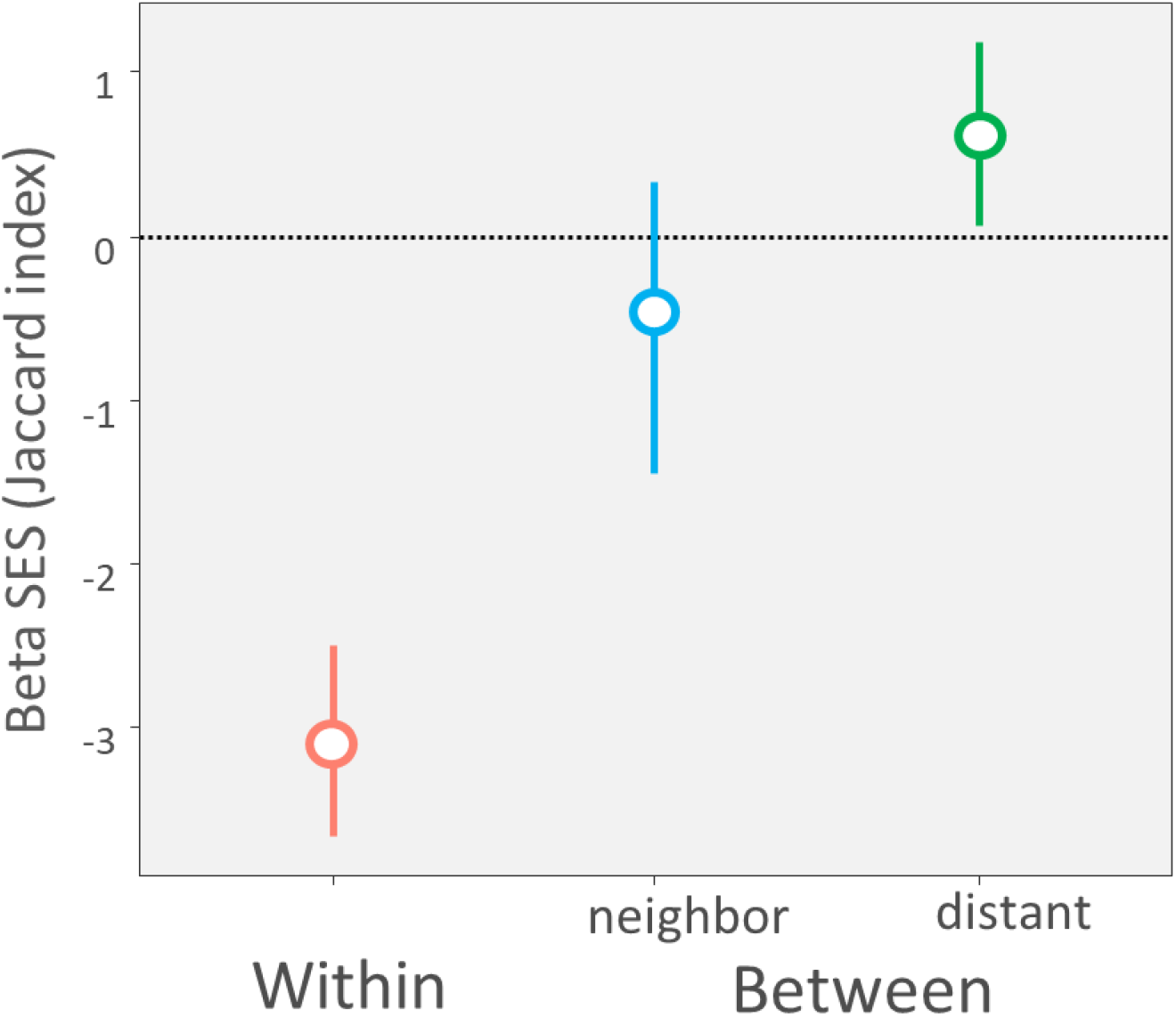
Standard effect size (SES) of the Jaccard index for comparison compositional dissimilarity within a same branch and between the adjacent and distant branches. Error bars indicate 99% confidence interval. The horizontal dotted line represents SES = 0, indicating a non-significant effect.

## Discussion

In the present study, we demonstrated the availability of eDNA metabarcoding for the investigation of fungal DNA assemblages and their spatial structure in river water. Our results show that the fungal DNA assemblages are more similar within branches than between. The results also show that the fungal DNA assemblages are more similar between spatially closer branches. Therefore, this study for the first time shows the spatial structure, especially compositional similarity, of fungal DNA assemblages across river branches.

### Fungal assemblages

River water contained both phylogenetically and functionally diverse fungal DNA. OTUs of terrestrial fungi such as the plant decomposer *Mycena* and the mycorrhizal fungi *Tuber* and Glomeromycota were detected, as well as the OTUs of aquatic fungi. These DNA may be released from the propagules derived from land. For example, Basidiomycota acting as plant decomposers often produce epigeous sporocarps and disperse their spores into air. Such airborne spores have the potential to enter the river system. On the other hand, species in the genus *Tuber* produce hypogenous sporocarps and do not disperse their spores into air. This indicates that the propagules and mycelial fragments in soil and plant materials (e.g., decomposing leaves) flow into the river via the surface current or with the direct addition of these substrates (Voronin, 2014). Recent studies that investigated the environmental fungal DNA of lakes and rivers also detected a significant amount of DNA for terrestrial fungi, as well as for aquatic fungi (Deiner et al., 2016, Khomich et al., 2017; LeBrun et al., 2018). Our results are consistent with the results of these previous studies.

The fungal OTU compositions in the studied landscape were more similar within the same branch than between branches. This may be because some fungal DNA, which entered upstream flowed downstream and was therefore detected both upstream and downstream in the branches. Indeed, recent evidence suggests that rivers work as a dispersal pathway of terrestrial fungi (Deiner et al., 2016; LeBrun et al., 2018). Other possibilities are that the same aquatic fungal species live across the branches, or that the same terrestrial fungal DNA enters the river at several different sites. However, there is currently no information in support of either of these possibilities. Further investigations into the composition and distribution of aquatic fungi along the river, the composition of fungal DNA flowing downstream, as well as the air and surface currents may help to clarify the mechanisms behind these results.

### The spatial structure of fungal DNA assemblages

Fungal DNA assemblages show geographical structures at a scale of around 15 km in the forest site. This means that fungal DNA assemblages are more similar between spatially closer branches than between distant ones. Such geographical structures have also been observed by Khomich et al. (2017); fungal DNA assemblages along latitudes and longitudes were investigated in a lake in the Scandinavia Peninsula. In the lake, the surrounding rivers brought and accumulated the DNA of terrestrial fungi from the surrounding areas (Khomich et al., 2017). Our study shows that such geographical structures can also be observed in streams, where water has a directional flow and is constantly replaced. The geographical structure of fungal DNA may be generated by both aquatic and terrestrial fungal DNA. For aquatic fungi, environmental differences (e.g. water chemical properties or current velocity) across branches or dispersal limitations can affect the fungal communities (Duarte et al., 2017), causing geographically closer branches to share more fungal species. Similarly, environmental factors such as vegetation type, soil chemical properties, and dispersal limitation contribute to the formation of similar terrestrial fungal communities in the same vicinity (Toljander et al., 2006; Kadowaki et al., 2014). The DNA of these communities flows into the nearby river, causing the DNA assemblages of geographically closer branches to resemble one another.

In the present study, no significant relationship between fungal DNA assemblage and the vegetation of the catchment area was detected. In a forest ecosystem, the types of vegetation present has a stronger effect on fungal communities compared to other environmental variables (Peay et al., 2013). Therefore, if the proportion of terrestrial fungal DNA is high, the fungal DNA assemblage may be explained by the vegetation present. There are two possible explanations for the lack of a relationship between the vegetative and fungal DNA assemblages. First, the fungal community may respond to factors other than vegetation types that were not measured in this study. For example, LeBrun et al., (2018) investigated fungal DNA assemblage in river water in an area with P enrichment problems and showed that the fungal DNA assemblage changes with P concentration. In our study sites, no such nutrient enrichment problems have been reported, although unmeasured factors and spatial distance should be considered in further studies. The second explanation is that the catchment area and distribution of fungal individuals of potential DNA origin did not correspond. For example, at a landscape scale with tens of kilometers, similar to our study site, fungal DNA would possibly flow or be blown from surrounding forests, including outside the catchment area. Otherwise, fungal DNA can be derived from areas closer to the river catchment area. There is limited information available on the dispersal ability of fungi. However, a previous study has reported that for terrestrial fungi, most spores fall within several meters of the sporocarps (Li et al., 2005; Galante et al., 2011). This indicates that spores of fungal species and individuals colonizing more than several meters apart from the river should have little chance to fall into river. Further studies are required to specify the range from which most fungal DNA is derived.

### Methodological limitations

In the present study, the ITS region was used as a genetic marker, but many OTUs remained unidentified. Compared to other terrestrial habitats, fewer genomic studies have been conducted in aquatic habitats. As a result, fewer ITS sequence records are available and new phylogenetic groups are expected to be found (Grossart et al., 2019; Khomich et al., 2018). Furthermore, for basal fungi living in an aquatic habitat, such as cystrids, more conservative regions such as large subunit (LSU) and small subunit (SSU) regions of rDNA are often used (Nilsson et al., 2019). However, few studies have investigated whether a difference in genetic markers affects the analytical results (Nilsson et al., 2019). Therefore, this may be an interesting topic for future studies to investigate. This database deficiency and handling of resulting unidentified OTUs as well as the selection of appropriate marker genes can be a key problem in fungal DNA metabarcoding.

## Conclusions

This is the first study to indicate that river water in a forest contains phylogenetically and functionally diverse fungal DNA that is derived from both land and aquatic regions. This study also suggests that assemblages of fungal DNA in river water also show a spatial structure. The results imply that by investigating fungal DNA assemblages in river water, information on both the land and aquatic fungal compositions of the catchment area can be accessed. Further study is required to clarify to what spatial extent of the fungal communities these DNA assemblages reflect, and what generates such a spatial structure of DNA assemblages.

## Acknowledgments

MiSeq sequencing was conducted in the Department of Environmental Solution Technology, Faculty of Science and Technology, Ryukoku University. We thank H. Yamanaka for supporting the MiSeq sequencing. This study received partial financial support from the Ministry of Education, Culture, Sports, Science, and Technology of Japan (MEXT) to SM (Grant No. 17K15199) and IK (18K11678) and the Environment Research and Technology Development Fund of Environmental Restoration and Conservation Agency (4-1602).

## Supporting information and Appendix

Table S1 List of all OTUs, their consensus sequence, and taxonomic and functional assignments.

Fig. S1 The relationship between the number of sequence reads and OTU numbers, i.e., rarefaction curves for the samples.

Fig. S2 Dissimilarity of the DNA assemblages among sites including Site I12 as revealed by nonmetric multidimensional scaling (NMDS) ordination (stress value = 0.1600). Numbers are consistent with site numbers in Fig. 1.

Fig. S3 PCA results of vegetation. PC1 and PC2 explain the variations in vegetation among the catchment area (see Fig. 1) for 86.9% and 11.2%, respectively.

Fig. S4 Phylum level proportions of the DNA assemblages for each sampling site.

Appendix 1. The results for additional analyses with different datasets.

