## Supporting Information for "Spatial structures of fungal DNA assemblages revealed with eDNA metabarcoding in a forest river network in western Japan"

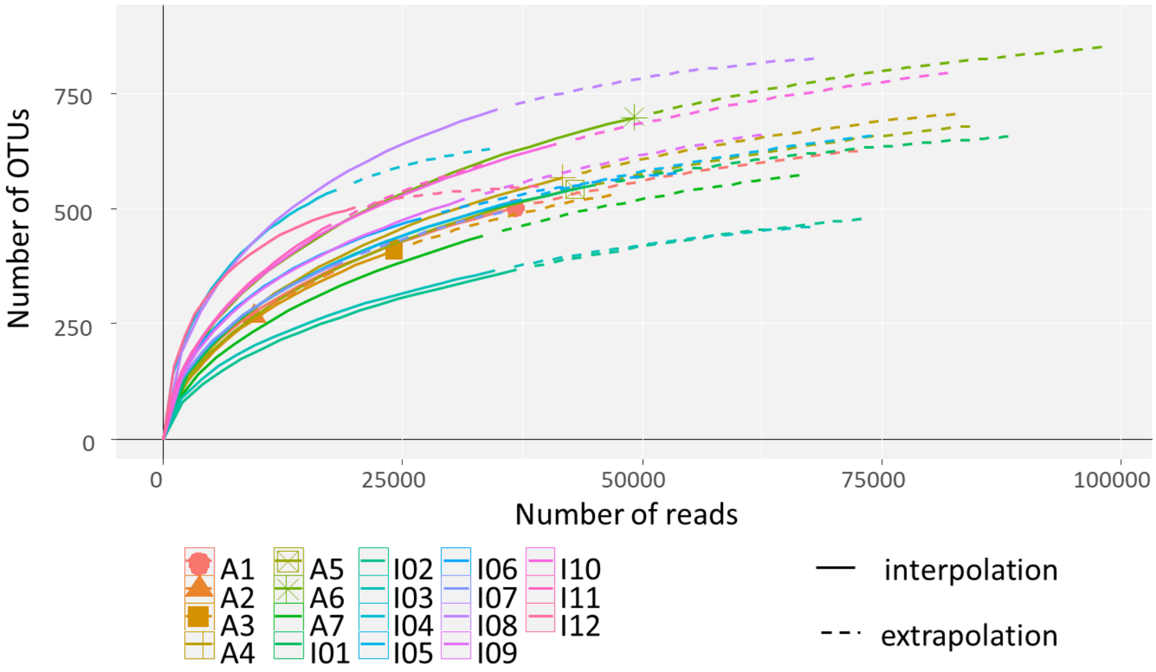


Fig. S1 The relationship between the number of sequence reads and OTU numbers, i.e., rarefaction curves for the samples.

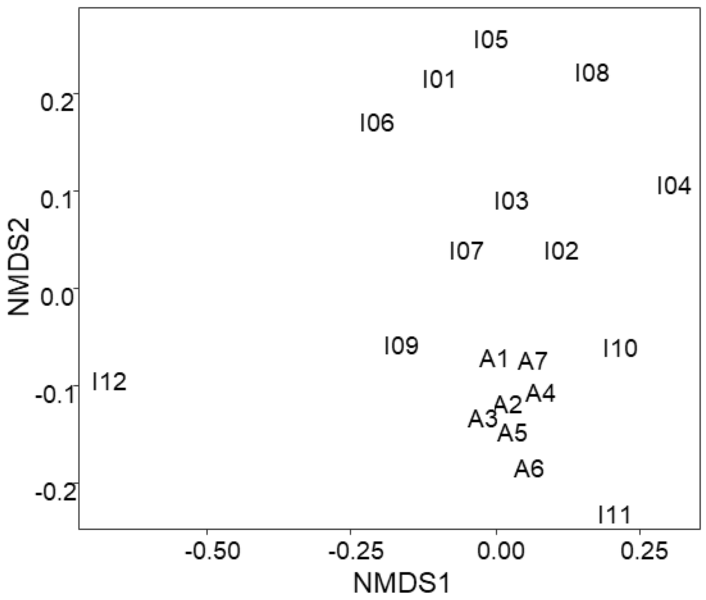


Fig. S2 Dissimilarity of the DNA assemblages among sites including Site I12 as revealed by nonmetric multidimensional scaling (NMDS) ordination (stress value = 0.1600). Numbers are consistent with site numbers in Fig. 1.

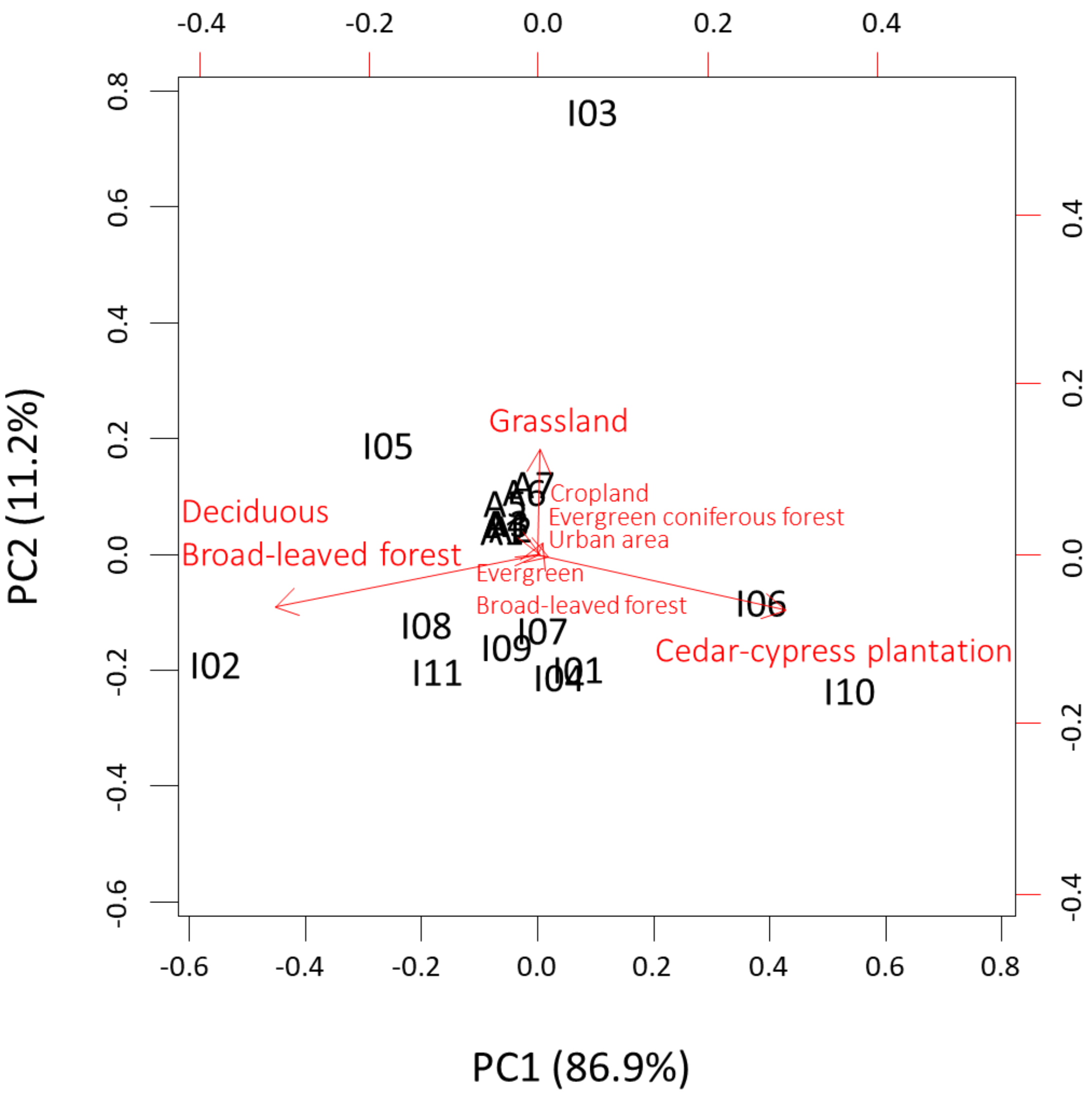


Fig. S3 PCA results of vegetation. PC1 and PC2 explain the variations in vegetation among the catchment area (see Fig. 1) for 86.9% and 11.2%, respectively.

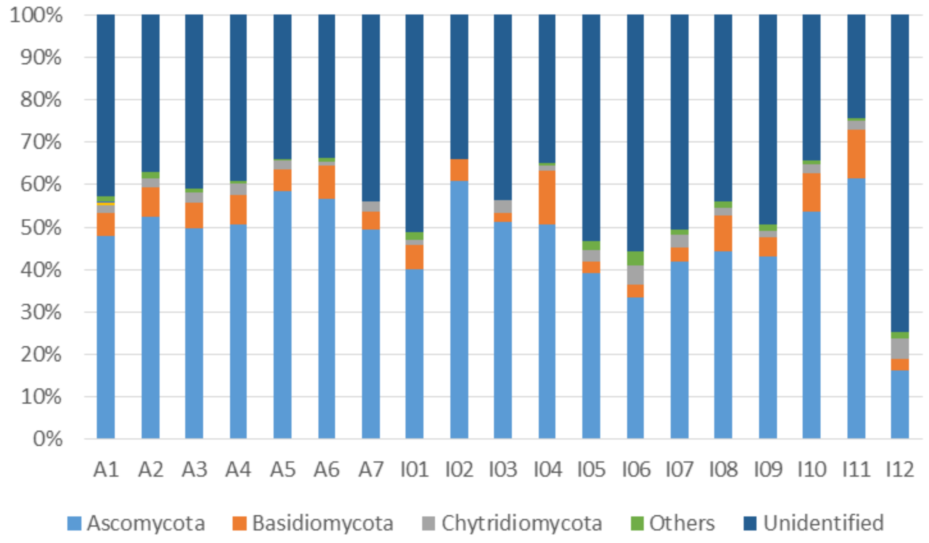


Fig. S4 Phylum level proportions of the DNA assemblages for each sampling site.
