## Appendix for "Spatial structures of fungal DNA assemblages revealed with eDNA metabarcoding in a forest river network in western Japan"

Appendix 1. The results for additional analyses with different datasets.

In the manuscript, results for the dataset employing presence/absence data for all OTUs, except for chimeric and non-fungal (but identified) OTUs, are shown. However, this dataset may include non-fungal OTUs. Therefore, the most conservative method of analysis is to include only those OTUs identified as fungus. In addition, some studies have treated a sequence read number as the proxy of abundance for each OTU, although there are known issues with the quantitative use of read numbers generated from amplicon sequencing (Amend *et al*., 2010; Elbrecht & Leese, 2015). Therefore, to test whether these differences in datasets can affect the analytical results, we conducted the same sets of analyses for the dataset including only fungal OTUs (Table S1) and using the sequence read number as a proxy of abundance for each OTU. The analytical procedure is the same as described in the manuscript except in the analyses considering the read number where the Bray-Curtis dissimilarity index was used instead of the Raup-Crick index. In addition, since the same randomization for presence/absence data cannot be conducted for non-binomial data, the pairwise comparison of SES values between and within branches (Fig. 3 in the main text) were not conducted on the dataset with the read number.

Both datasets yielded the same results as in the manuscript; the dissimilarity of OTU composition among sites correlated with the latitude and longitude (Fig. A1-1). The OTU composition was more similar within the same branches than between branches (Fig A1-1 and Fig A1-1) and only the PCNM1 vector significantly related to the difference in OTU composition among sites (Table A1-1). These results indicate that the result shown in the manuscript is not vulnerable to the dataset choice.

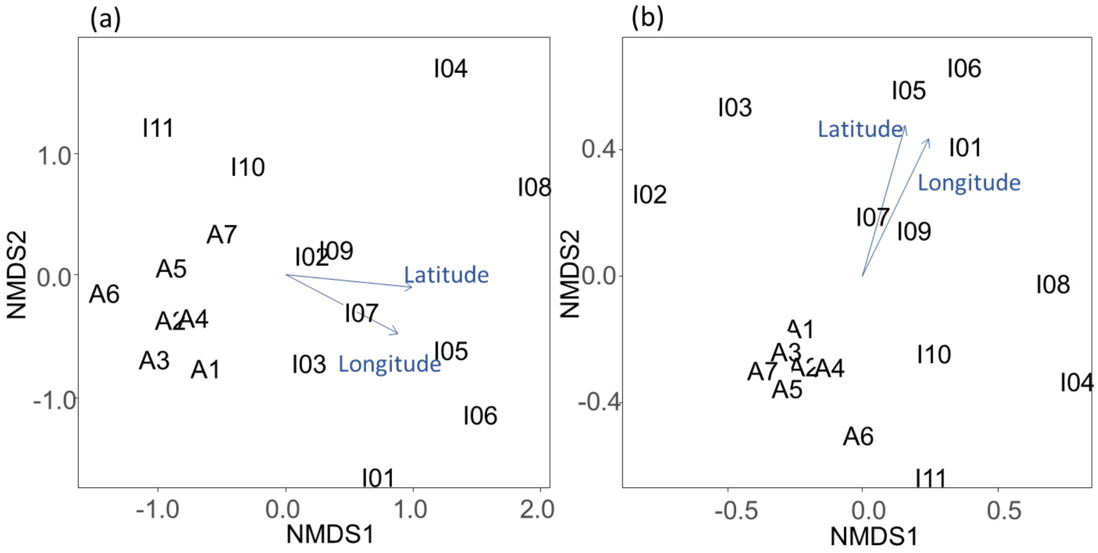


Fig. A1-1 Dissimilarity of the DNA assemblages among sites as revealed by nonmetric multidimensional scaling (NMDS) ordination for (a) datasets with only fungal OTUs (stress value = 0.159) and (b) datasets using the read number (stress value = 0.153). Numbers are consistent with site numbers in Fig. 1 in the main text. The ordinations were significantly correlated with latitude and longitude for both datasets (‘envfit’ function; latitude, (a) $r^2 = 0.749$, $P = 0.0002$, (b) $r^2 = 0.684$, $P = 0.0006$; longitude, (a) $r^2 = 0.7931$, $P = 0.0002$, (b) $r^2 = 0.802$, $P = 0.0002$).

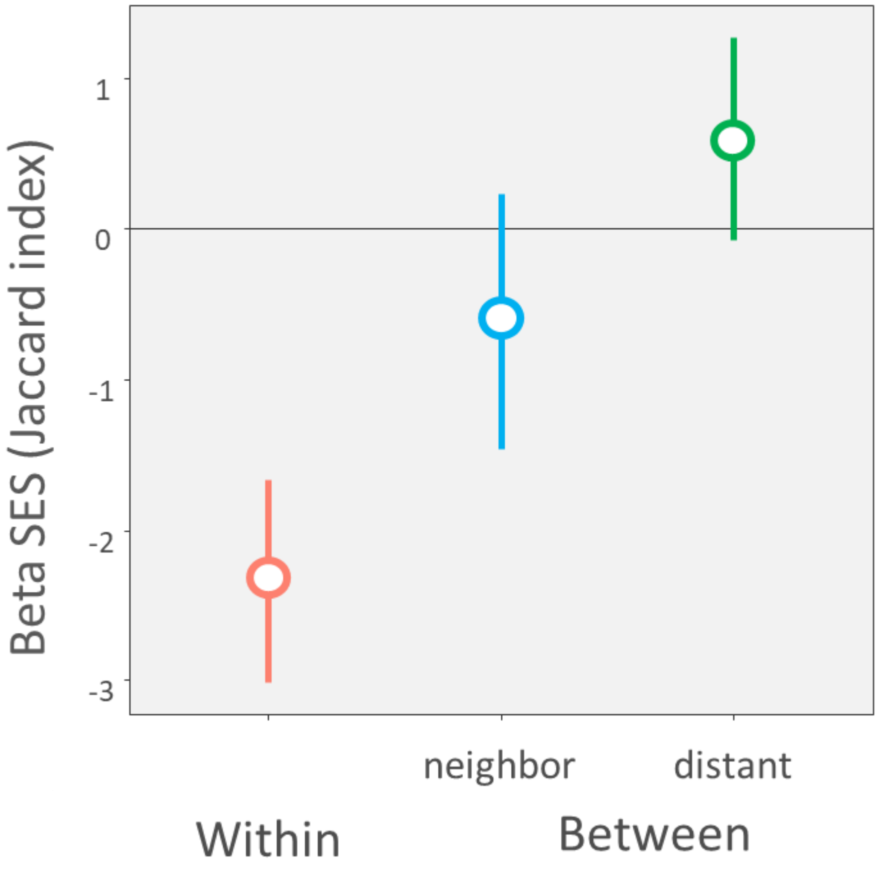


Fig. A1-2 Standard effect size (SES) of the Jaccard index for comparison compositional dissimilarity within a same branch and between the adjacent and distant branches. Error bars indicate 99% confidence interval. The horizontal dotted line represents SES = 0, indicating a non-significant effect.

Table A1-1 PERMANOVA results for the composition of DNA assemblages

| | Degree of freedom | Sums of Squares | F Model | $R^2$ | P value |
|---|---|---|---|---|---|
| (a) Only fungal OTUs included | | | | | |
| null | 10 | 1.900 | | | |
| Elevation | 1 | 0.334 | 1.918 | 0.176 | 0.222 |
| Catchment area | 1 | 0.173 | 0.900 | 0.091 | 0.494 |
| Vegetation PC1 | 1 | 0.167 | 0.866 | 0.088 | 0.485 |
| Vegetation PC2 | 1 | -0.106 | -0.476 | -0.056 | 0.954 |
| PCNM1 | 1 | 0.794 | 6.459 | 0.418 | 0.007 |
| PCNM2 | 1 | 0.017 | 0.080 | 0.009 | 0.781 |
| (b) When read number considered | | | | | |
| null | 10 | 1.194 | | | |
| Elevation | 1 | 0.452 | 2.931 | 0.246 | 0.126 |
| Catchment area | 1 | 0.269 | 1.540 | 0.146 | 0.335 |
| Vegetation PC1 | 1 | 0.159 | 0.849 | 0.086 | 0.501 |
| Vegetation PC2 | 1 | -0.015 | -0.075 | -0.008 | 0.863 |
| PCNM1 | 1 | 0.766 | 6.416 | 0.416 | 0.012 |
| PCNM2 | 1 | -0.088 | -0.411 | -0.048 | 0.905 |
